# Cell-type specific eQTL of primary melanocytes facilitates identification of melanoma susceptibility genes

**DOI:** 10.1101/231423

**Authors:** Tongwu Zhang, Jiyeon Choi, Michael A. Kovacs, Jianxin Shi, Mai Xu, NISC Comparative Sequencing Program, Melanoma, Meta-Analysis Consortium, Alisa M. Goldstein, Mark M. Iles, David Duffy, Stuart MacGregor, Laufey T. Amundadottir, Matthew H. Law, Stacie K. Loftus, William J. Pavan, Kevin M. Brown

## Abstract

Most expression quantitative trait loci (eQTL) studies to date have been performed in heterogeneous tissues as opposed to specific cell types. To better understand the cell-type specific regulatory landscape of human melanocytes, which give rise to melanoma but account for <5% of typical human skin biopsies, we performed an eQTL analysis in primary melanocyte cultures from 106 newborn males. We identified 597,335 *cis*-eQTL SNPs prior to LD-pruning and 4,997 eGenes (FDR<0.05), which are higher numbers than in any GTEx tissue type with a similar sample size. Melanocyte eQTLs differed considerably from those identified in the 44 GTEx tissues, including skin. Over a third of melanocyte eGenes, including key genes in melanin synthesis pathways, were not observed to be eGenes in two types of GTEx skin tissues or TCGA melanoma samples. The melanocyte dataset also identified cell-type specific *trans-*eQTLs with a pigmentation-associated SNP for four genes, likely through its *cis*-regulation of *IRF4*, encoding a transcription factor implicated in human pigmentation phenotypes. Melanocyte eQTLs are enriched in *cis*-regulatory signatures found in melanocytes as well as melanoma-associated variants identified through genome-wide association studies (GWAS). Co-localization of melanoma GWAS variants and eQTLs from melanocyte and skin eQTL datasets identified candidate melanoma susceptibility genes for six known GWAS loci including unique genes identified by the melanocyte dataset. Further, a transcriptome-wide association study using published melanoma GWAS data uncovered four new loci, where imputed expression levels of five genes (*ZFP90, HEBP1, MSC, CBWD1*, and *RP11-383H13.1*) were associated with melanoma at genome-wide significant *P*-values. Our data highlight the utility of lineage-specific eQTL resources for annotating GWAS findings and present a robust database for genomic research of melanoma risk and melanocyte biology.

## INTRODUCTION

Expression quantitative trait locus (eQTL) analysis is a powerful method to study gene expression and regulatory profiles in human populations. Early studies mainly focused on eQTLs for whole blood or blood-derived cells due to sample accessibility (Stranger et al. 2007; Pickrell et al. 2010), and more recently, numerous eQTL datasets derived from normal human tissues have been made publicly available. Perhaps most notable are those from the Genotype-Tissue Expression (GTEx) (The GTEx Consortium 2015) project representing >44 tissue types of hundreds of post-mortem donors. These studies have collectively emphasized the cell-type specific nature of eQTLs, where 29-80% of eQTLs are cell-type specific (Dimas et al. 2009; Nica et al. 2011; Fairfax et al. 2012; The GTEx Consortium 2015). While eQTLs from normal tissues provide valuable insights, tissues are constituted of multiple distinct cell types with specific gene regulatory profiles as exemplified by eQTLs of different blood-isolated cell types (Fairfax et al. 2012). Moreover, the collection and sampling process of tissue samples from organs does not allow precise control over cell representation, adding a major source of biological variability in addition to other technical variation (McCall et al. 2016). However, other than for immune cells (Kim-Hellmuth et al. 2017) or induced Pluripotent Stem Cells (iPSC) (Kilpinen et al. 2017), eQTL datasets representing single primary cell types and direct comparison of these to the tissue type of origin have been lacking.

eQTLs may be particularly useful for annotating variants associated with complex traits, as such variants are likely enriched for eQTLs (Nicolae et al. 2010). A recent study suggested that two-thirds of candidate common trait susceptibility genes identified as eQTLs are not the nearest genes to the GWAS lead SNPs, highlighting the utility of this approach in annotating GWAS loci (Zhu et al. 2016). Importantly, GWAS variants are enriched in eQTLs in a tissue-specific manner. For instance, whole blood eQTLs are enriched with autoimmune disorder-associated SNPs but not with GWAS SNPs for bipolar disease or type 2 diabetes (The GTEx Consortium 2015). These findings highlight the importance of using eQTL datasets from relevant cell types when following up GWAS loci for a specific disease. In addition to providing functional insights for known GWAS loci, eQTL data may be useful for identification of novel trait-associated loci via imputation of genotype-correlated gene expression levels into GWAS datasets (Gamazon et al. 2015; Gusev et al. 2016). Such approaches, usually referred to as transcriptome-wide association studies (TWAS) enable assignments of potentially disease-associated loci via estimations of their genetically regulated expression.

GWAS for melanoma risk, nevus count, and multiple pigmentation traits have identified numerous associated genetic loci (Stokowski et al. 2007; Sulem et al. 2007; Brown et al. 2008; Gudbjartsson et al. 2008; Han et al. 2008; Sulem et al. 2008; Bishop et al. 2009; Falchi et al. 2009; Nan et al. 2009; Duffy et al. 2010; Eriksson et al. 2010; Amos et al. 2011; Barrett et al. 2011; Macgregor et al. 2011; Nan et al. 2011; Candille et al. 2012; Zhang et al. 2013; Jacobs et al. 2015; Law et al. 2015; Liu et al. 2015; Hysi et al. 2018; Visconti et al. 2018), with melanoma GWAS alone identifying 20 regions associated with risk. Trait-associated variation explaining many of these loci could reasonably be expected to be reflected in the biology of the melanocyte, the pigment producing cell in human skin and the cellular origin of melanoma. Melanocytes are the cells in the skin that function to produce the melanin pigments, eumelanin and pheomelanin, in response to neuroendocrine signals and UV-exposure (Costin and Hearing 2007). These melanin pigments are contained in lysosome-related organelles called melanosomes, are shuttled to the melanocyte dendrites, and transferred to neighboring keratinocytes thus protecting skin from UV radiation (Sitaram and Marks 2012). The process of pigmentation is complex and multigenic, and it is regulated by genes with diverse cellular functions including those within MAPK, PI3K, Wnt-/beta catenin signaling pathways (Liu et al. 2014) as well as those involved in lysosome-related functions and vesicular trafficking (Sitaram and Marks 2012).

While several skin-related eQTL datasets are available, the largest ones (GTEx (The GTEx Consortium 2015), MuTHER (Nica et al. 2011), EUROBATS (Buil et al. 2015)) are derived from heterogeneous skin tissues, of which melanocytes only represent a small fraction. The Cancer Genome Atlas Project (TCGA) also offers a considerable set of tumor tissue expression data accompanied by genotype information providing a platform for tumor-type relevant eQTL data including melanoma (https://cancergenome.nih.gov/), but these tumor tissues contain a high burden of somatic aberrations, are heterogeneous and may reflect multiple disease subtypes, and may not represent the underlying biology associated with cancer risk and/or pigmentation.

Given these limitations, we took advantage of the accessibility of primary melanocytes obtained from foreskin tissues and built a cell-type specific eQTL dataset to study the lineage-specific regulatory function of melanoma- and pigmentation-associated common variants.

## RESULTS

### Melanocyte eQTLs are distinct from those of other tissue types

In order to create a melanocyte-specific eQTL resource, we obtained primary melanocyte cultures isolated from foreskin of 106 healthy newborn males predominantly of European descent (**Supplemental Table 1**). We then cultured all 106 lines following a uniform procedure to harvest RNA and DNA, for RNA sequencing and genotyping, respectively (**see Methods**). Given the relatively small size of our sample set, we initially focused our analysis on local eQTLs (*cis*-eQTL), where we assessed the association between expression of each gene with common variants within +/−1Mb of transcription start sites (TSS), following the best practices from the GTEx project (see **Methods**). In all, we identified 4,997 “eGenes” (genes exhibiting association with genotypes of at least one SNP at FDR < 0.05; **Supplemental Table 2**) and 597,335 genome-wide “significant eQTLs” (unique SNP-gene pairs showing FDR < 0.05; SNPs were not LD-pruned), which are higher numbers than any GTEx tissue type of similar sample size (**Supplemental Table 3**). Melanocyte eGenes were enriched with Gene Ontology (GO) terms including metabolic process, mitochondrial translation, biosynthetic process, catalytic activity and ion-binding, as well as lysosome and metabolic pathways (**Supplemental Table 4**). Further, melanocyte eGenes included 46% of genes categorized with GO terms as containing “melanin” (*OCA2, TRPC1, CTNS, DCT, MCHR1, SLC45A2, TYR, BCL2, WNT5A, MC1R*, and *MYO5A*) (http://amigo.geneontology.org), and 20% of curated pigmentation genes (based on human and mouse phenotype, OMIM, MGI) such as *IRF4, TRPM1*, and *MC1R* (**Supplemental Table 5-6**), reflecting pigmentation related biology of melanocytes.

Direct comparison of significant melanocyte eQTLs with 44 GTEx tissue types indicated that the shared eQTL proportion (*π*_1_) between melanocytes and each of GTEx tissue types was 0.74 (vs. transformed fibroblasts) or lower, suggesting relatively low levels of sharing even with two types of skin samples (*π*_1_ = 0.67 with Skin_Sun_Exposed, and 0.58 with Skin_Not_Sun_Exposed; **Fig. 1**). This contrasts with the considerably higher levels of sharing between the two types of skin samples (*π*_1_ = 0.91) or among brain tissues (average *π*_1_=0.87) in GTEx. We further focused the comparison of our melanocyte dataset to three tissue types that are directly relevant to melanoma and pigmentation phenotypes: the two above-mentioned GTEx skin types, as well as skin cutaneous melanomas (SKCM) collected through TCGA (adding an adjustment for local DNA copy number). Collectively, these four eQTL datasets identified 12,136 eGenes, with 382 eGenes shared among all four datasets. Notably, 1,801 eGenes (36% of melanocyte eGenes) were entirely private to melanocytes, and a total of 6,187 eGenes (51% of eGenes from all four datasets) were specific to only one of four datasets (**Supplemental Fig. S1; Supplemental Table 2**). eGenes from these four datasets collectively accounted for 150 of 379 (40%) curated pigmentation genes, with the majority specific to one dataset (**Supplemental Fig. S2**).

**Figure 1.**
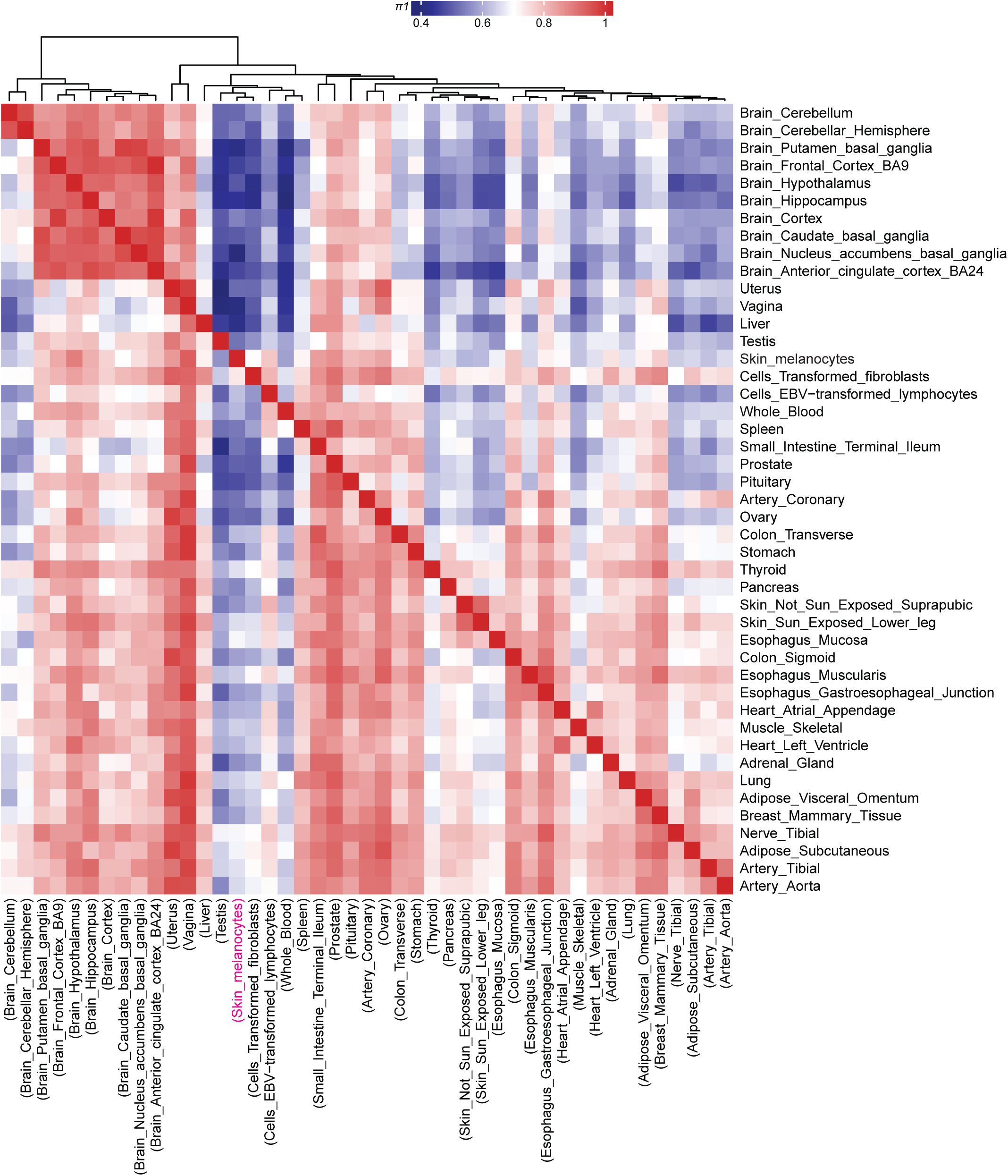
Melanocyte eQTLs display a distinct pattern from those of 44 GTEx tissue types. Dendrogram and heatmap presenting the sharing of eQTLs between human primary melanocytes and 44 other GTEx tissue types. Pairwise *π*_1_ statistics were calculated from single-tissue eQTL discoveries in each tissue using all the genome-wide significant eQTL SNP-gene pairs. *π*_1_ is only calculated when the gene is expressed and testable both in discovery (columns) and replication (rows) tissues. Higher *π*_1_ values indicate an increased replication of eQTLs between two tissue types. *π*_1_ values range between ~0.41 and 1, and are color-coded from blue (low sharing) to red (high sharing). Tissues are clustered using the Spearman correlation of *π*_1_ values. Note that *π*_1_ values are not symmetrical, since each entry in row *i* (replication tissue) and column *j* (discovery tissue) is an estimate of *π*_1_ = *Pr* (eQTL in tissue *i* given an eQTL in tissue *j*). Discovery tissue names are shown in parenthesis on the bottom. The position of the skin melanocyte eQTL dataset from the discovery tissues is shown in pink.

### Melanocyte eQTLs are enriched in *cis*-regulatory signatures and supported by allelic imbalance

We next sought to determine whether melanocyte eQTLs were corroborated by allelic imbalance variants in heterozygous individuals from the same dataset. To determine genome-wide allele-specific expression (ASE), we performed binomial tests at the single sample level, identifying 48,038 unique allelic imbalance variants (FDR < 0.05 or effect size > 0.15; **Supplemental Table 7**). Of these unique variants 38.6% (18,532 of 48,038 variants) were in the coding region of significant melanocyte eGenes, demonstrating an enrichment of ASE in melanocyte eGenes (Fisher’s exact test *P* = 2.34 × 10^−73^, Odds Ratio = 1.82; **Supplemental Table 7**). Further, the average allelic effects of 48,038 ASE variants from all the heterozygous individuals were significantly larger in the eGene group (Wilcoxon signed rank test *P* = 1.67 × 10^−34^; average |Mean AE| = 0.046 for eGenes vs. 0.035 for non-eGenes; effect size = 0.115; **Supplemental Fig. S3A**). Similarly, the proportions of heterozygous individuals displaying allelic imbalance at each locus was significantly higher in the eGene group (Wilcoxon signed rank test *P* = 1.27 × 10^−81^; mean % = 13.4 for eGenes vs. 8.4 for non-eGenes; effect size = 0.195; **Supplemental Fig. S3B**).

We then further examined if melanocyte eQTLs were enriched within epigenetic signatures marking melanocyte *cis*-regulatory elements. We specifically examined regions of open chromatin (marked by DNaseI hypersensitivity sites; DHS)), as well as promoter and enhancer histone marks (H3K27ac, H3K4me1, and H3K4me3) generated from primary cultured human melanocytes by the ENCODE and Epigenome Roadmap Projects (www.encodeproject.org; www.roadmapepigenomics.org) (ENCODE Project Consortium 2012) (Roadmap Epigenomics et al. 2015). Indeed, higher proportions of melanocyte eQTL SNPs were localized to melanocyte DHS, H3K27ac, H3K4me1, and H3K4me3 peaks compared to all tested SNPs (i.e. *cis-SNPs* +/− 1Mb of TSS of all the genes tested for eQTL) (**Supplemental Fig. S4A**). Enrichment of melanocyte eQTL SNPs for each of the melanocyte *cis*-regulatory signatures was statistically significant (*P* < 1 × 10^−4^, 10,000 permutations; 1.81 to 5.48-fold) (**Table 1**) and mostly more pronounced than that observed in GTEx skin tissues or melanoma tumors (**Supplemental Fig. 4B; Table 1**). Further, melanocyte eQTL SNPs were also enriched (permutation test *P* < 1 × 10^−4^) upstream of genes (within 1-5 kb of the TSS; ~2.5 fold), as well as in gene promoters, 5’ UTRs, exonic regions, first exons, first introns, introns, intron-exon boundaries (+/− 200bp encompassing exon splice regions), and 3’ UTRs, but not in intergenic regions or annotated lincRNA regions (**Supplemental Fig. S4**). Consistent with this enrichment, most of the melanocyte eQTL SNPs were centered within +/− 250kb of transcription start sites (TSS) (**Supplemental Fig. S5**).

**Table 1.** Enrichment of eQTL SNPs in melanocyte *cis*-regulatory signatures.

| Epigenetic Mark |  | DHS | H3K27ac | H3K4Me1 | H3K4Me3 |
| --- | --- | --- | --- | --- | --- |
| **Melanocyte eQTLs** | Fold enrichment | 1.81 | 5.48 | 1.99 | 3.34 |
|  | P-value | <0.0001 | <0.0001 | <0.0001 | <0.0001 |
| **TCGA SKCM eQTLs** | Fold enrichment | 1.72 | 2.69 | 1.86 | 3.44 |
|  | P-value | <0.0001 | <0.0001 | <0.0001 | <0.0001 |
| **Skin Not Sun Exposed eQTLs** | Fold enrichment | 1.78 | 2.77 | 1.92 | 3.5 |
|  | P-value | <0.0001 | <0.0001 | <0.0001 | <0.0001 |
| **Skin Sun Exposed eQTLs** | Fold enrichment | 1.75 | 2.66 | 1.88 | 3.29 |
|  | P-value | <0.0001 | <0.0001 | <0.0001 | <0.0001 |

Epigenetic Mark: DNaseI hypersensitivity sites (DHS) and gene regulatory histone marks (H3K27ac, H3K4me1, and H3K4me3) of primary melanocytes (Epigenome Roadmap database)
Fold enrichment: mean fold enrichment of eQTL SNPs over control SNP sets (with similar distribution of MAF and LD) from 10,000 permutations that are overlapping with each epigenetic mark.
P-value: see **Methods** section

### Melanoma GWAS signal is enriched in melanocyte-specific genes and eQTLs

Next, we sought to determine if melanocyte eQTLs were enriched with common risk variants from the most recent melanoma GWAS meta-analysis (Law et al. 2015). Quantile-quantile plot demonstrated an enrichment of significant GWAS *P*-values for eQTL SNPs compared to non-eQTL SNPs (**Fig. 2A**), which was the most pronounced in melanocyte eQTL (estimated Lambda = 1.51) compared to three related tissue types as well as all the other GTEx tissue types (**Supplemental Fig. S6**).

**Figure 2.**
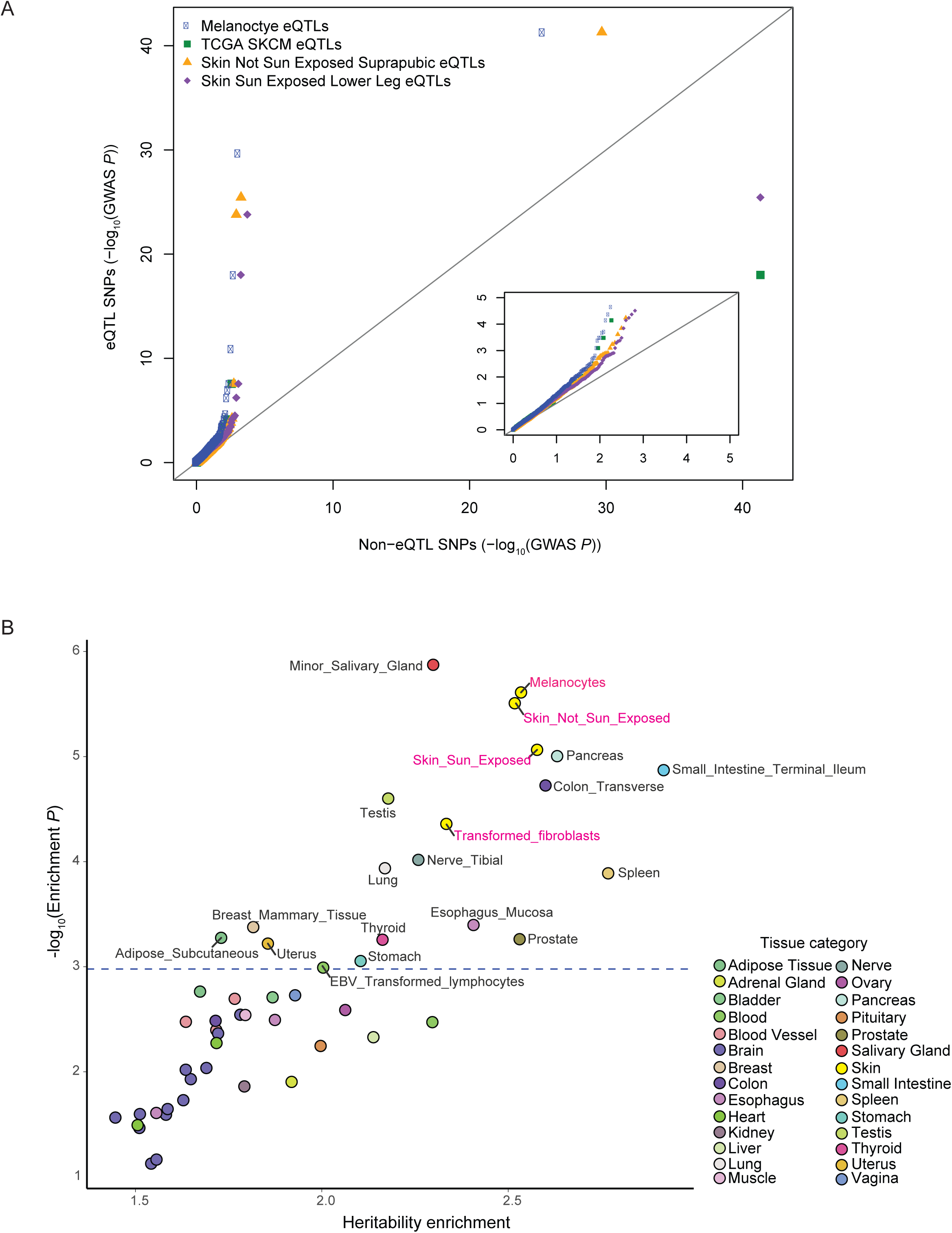
Melanoma GWAS signal is enriched in melanocyte-specific genes and eQTLs. (*A*) QQ plot presents melanoma GWAS LD-pruned *P*-values of significant eQTL SNPs versus non-eQTL SNPs for the melanocyte dataset compared to those for sun-exposed skin, non-sun-exposed skin, and melanoma tumors. SNPs were classified as eQTL SNPs if they were significant eQTLs or in strong LD (*r*^2^>0.8) with an eQTL SNP (eQTL SNPs threshold: FDR < 0.05) in each dataset. The inset displays a zoomed-in view of a lower −log10 GWAS *P*-value range (0-5 range for X- and Y-axes). (*B*) Melanoma heritability enrichment levels and *P*-values in top 4000 tissue-specific genes from LD score regression analysis are displayed. The dashed horizontal line marks FDR = 0.05 on the Y-axis. Names of significantly enriched individual tissue types are shown next to the data points, and the others are color-coded based on GTEx tissue category. Tissue types from the “Skin” category including melanocytes are highlighted in pink.

To further assess the enrichment of melanoma heritability in melanocyte-specific expressed genes, we performed LD score regression analysis (Finucane et al. 2015). The results indicated that partitioned melanoma heritability was significantly enriched (2.54 fold; *P* = 2.45 × 10^−6^) in melanocyte-specific genes (top 4,000 genes compared to 47 GTEx tissue types), as well as in those of three “skin” category GTEx tissue types, albeit to a lesser degree (*P* = 3.11 × 10^−6^, 8.62 × 10^−6^, and 4.37 × 10^−5^, with 2.52, 2.58, and 2.34 fold for not sun-exposed skin, sun-exposed skin, and transformed fibroblasts, respectively) (**Fig. 2B; Supplemental Table 8; Supplemental Fig. S7**).

### A functional pigmentation SNP at the *IRF4* locus is a significant *trans*-eQTL for four genes in melanocytes

While the modest size of this dataset limits power, we also performed *trans-eQTL* analyses for the SNPs that are located over 5Mb away from the TSS of each gene or on a different chromosome. In all, we identified 15 genome-wide significant *trans*-eQTL genes (excluding genes of mappability < 0.8 or overlapping low complexity regions; **Supplemental Table 9**). Of these, eight *trans*-eQTL SNPs were also *cis*-eQTLs for local genes within 1Mb. Notably, rs12203592 (Chr6:396321), among these, is a genome-wide significant *trans*-eQTL SNP for four different genes on four separate chromosomes (*TMEM140, MIR3681HG, PLA1A*, and *NEO1*) and is also the strongest *cis*-eQTL SNP for the *IRF4* gene (P = 7.9 × 10^−16^, slope = − 1.14), which encodes the transcription factor, interferon regulatory factor 4. All four genes displayed the same direction of allelic gene expression as *IRF4* levels relative to rs12203592 (**Fig. 3**). rs12203592 has previously been associated with human pigmentation phenotypes (Han et al. 2008). This variant was also shown to be a functional SNP mediating transcription of *IRF4* in melanocytes via C allele-preferential binding of the transcription factor, TFAP2, by collaborating with melanocyte-lineage specific transcription factor, MITF, in turn activating the melanin synthesis enzyme, *TYR*. The rs12203592-C allele (prevalent in African populations) is correlated with high *IRF4* levels in melanocytes, validating the findings observed in a smaller sample set (Praetorius et al. 2013). Expression correlation analyses in melanocytes indicated that expression levels of *TMEM140, MIR3681HG, PLA1A*, and *NEO1* are significantly correlated with those of *IRF4* in the same direction as shown by *trans*-eQTL (Pearson *r* = 0.54, 0.65, 0.53, and 0.58; *P* = 2.67 × 10^−9^, 5.34 × 10^−14^, 4.28 × 10^−9^, and 6.00 × 10^−11^, respectively; **Supplemental Fig. S8**). To assess if *IRF4* expression levels mediate the observed *trans*-eQTL effect for these four genes, we performed mediation analyses in three different ways (**Supplemental Material**). Briefly, we used regression of *trans*-eQTL gene levels against rs12203592 either by taking the residuals after accounting for *IRF4* levels or using *IRF4* levels as a covariate. The results indicated considerable increases in association *P*-values of the residuals for all four genes, but no significant change with or without *IRF4* as a covariate (**Supplemental Table 10**). We also applied a recently published Genomic Mediation analysis with Adaptive Confounding adjustment (GMAC) (Yang et al. 2017) to 455 eSNP - *cis*-eGene - *trans-gene* trios (*trans*-eQTL cutoff: *P* < 1 × 10^−5^), 84 of which include rs12203592. A total of 121 trios displayed a suggestive mediation (*P* < 0.05), and 32 of them were by *IRF4 cis*-eQTL including those with *TMEM140* and *NEO1* (**Supplemental Table 11**). In contrast, another *cis*-eQTL gene, *RPS14*, sharing two SNPs with three *trans*-eQTL genes (**Supplemental Table 9**), did not show suggestive mediation (**Supplemental Table 11**). These results are consistent with *IRF4* expression levels mediating at least part of the observed *trans*-eQTL effect. We then sought to determine if IRF4 is predicted to bind to the genomic regions encompassing the rs12203592 *trans*-eQTL genes. Sequence motif enrichment analyses indicated that IRF4 binding motifs were enriched in the genomic regions of *TMEM140, MIR3681HG, PLA1A*, and *NEO1* (+/− 2 kb of gene boundary; *P* = 1.52 × 10^−2^; Supplemental Table 12), as well as in above-mentioned 84 *trans*-eQTL genes (*P* = 7.25 × 10^−26^). Together our data suggest a melanocyte-specific *trans*-eQTL network potentially regulated by the transcription factor, IRF4.

**Figure 3.**
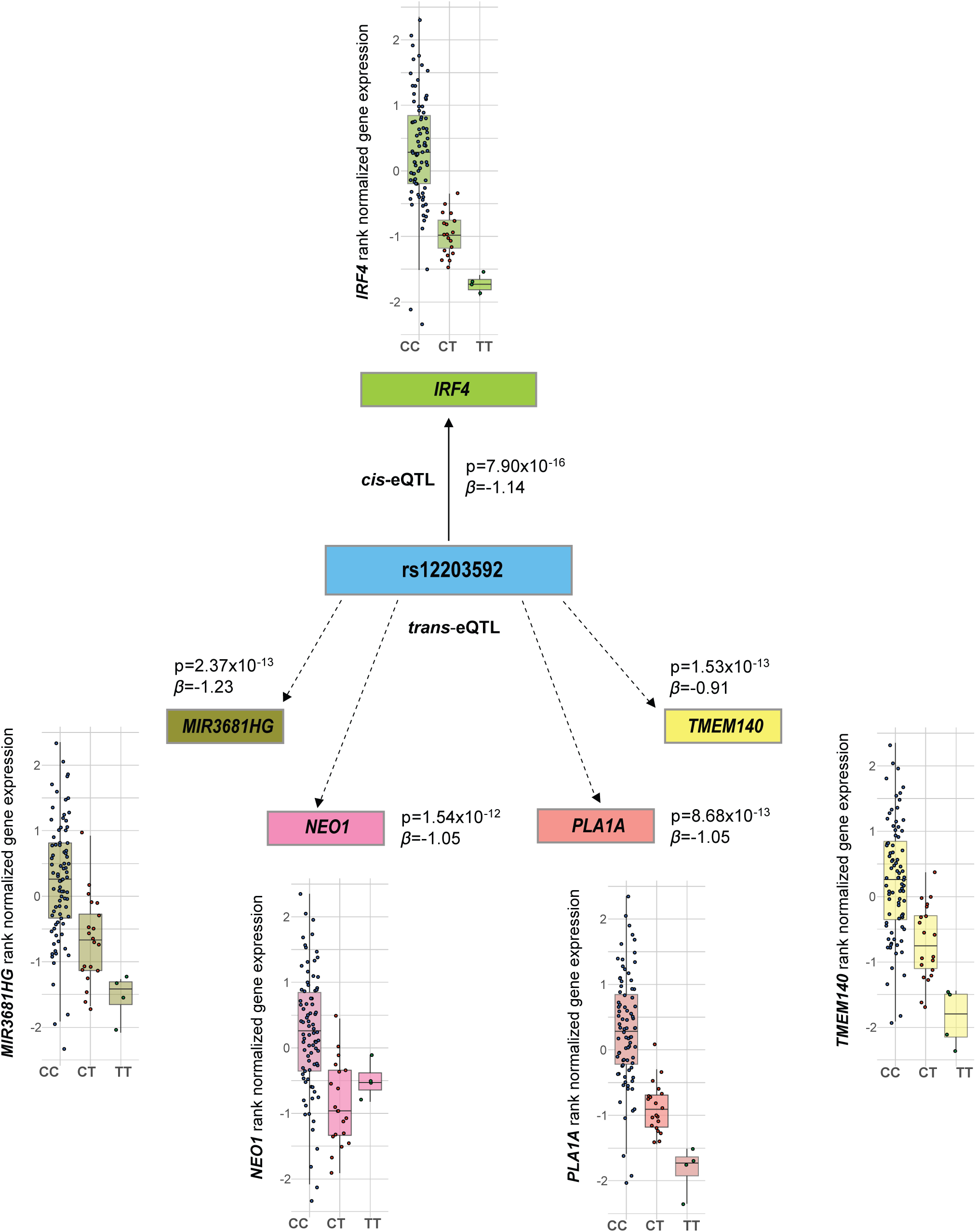
The pigmentation trait-associated variant, rs12203592, in *IRF4* is a *trans*-eQTL for four genes in melanocytes. *cis-* or *trans-*eQTL *P*-values and effect sizes (β) are shown between rs12203592 and *IRF4* or rs12203592 and four genome-wide significant *trans*-eQTL genes (*TMEM140, MIR3681HG, PLA1A*, and *NEO1*). β values are shown relative to alternative alleles, T. Boxplots display gene expression levels based on rs12203592 genotypes (CC, CT, and TT).

### Melanocyte eQTLs identified candidate melanoma susceptibility genes from GWAS loci

To assess colocalization of causal variants for melanoma GWAS and melanocyte eQTL, we applied the previously described eCAVIAR methodology (Hormozdiari et al. 2016). At a colocalization posterior probability (CLPP) cutoff of 1%, 5 of 20 known melanoma loci displayed colocalization of GWAS and melanocyte eQTL signal, with colocalization of eQTL signal for nine genes overall (**Table 2; Fig. 4**). The same analysis with two GTEx skin datasets observed colocalization at combined four loci and 21 genes (**Supplemental Table 13**). The union of all three datasets totaled 29 genes from six loci, indicating that these eQTL datasets complement each other rather than being redundant. Consistent with a previous report (The GTEx Consortium 2017), only 66% (4 of 6 loci) but not all of melanoma GWAS signal colocalized with the nearest expressed gene in one or more of the three datasets. Importantly, melanocyte eQTL (but not the skin datasets) validated *PARP1* as a target gene on the locus at Chr1q42.12, which was previously characterized as a melanoma susceptibility gene displaying melanocyte lineage-specific function (Choi et al. 2017) (**Fig. 4**). Melanocyte eQTL also uniquely identified a known pigmentation gene, *SLC45A2*, on the locus at 5p13.2 as a target gene, reflecting a melanin synthesis pathway uniquely captured in melanocyte eQTL. Consistent with previous findings, eCAVIAR colocalization was observed for multiple genes in most of the loci, and genes with the highest CLPP scores from different eQTL datasets did not overlap for a given melanoma locus. In addition, we also performed eCAVIAR analyses for GWAS of melanoma-associated traits (number of melanocytic nevi, skin pigmentation, ease of tanning, and hair color), and identified target genes from two of four nevus count loci and six of 11 pigmentation loci using melanocyte and skin eQTL datasets (**Supplemental Material; Supplemental Table 14-16**).

**Table 2.** Colocalization of melanoma GWAS and melanocyte eQTL signal^*^.

| Melanoma GWAS Locus | Gene | SNP ID | GWAS *P*-value | CLPP | Melanoma GWAS Lead SNP | GWAS *P*-value | *r*<sup>2</sup> | Nearest Expressed Gene |
| --- | --- | --- | --- | --- | --- | --- | --- | --- |
| **1q21.3** | **ARNT** | **rs12410869** | **5.21E-13** | **0.07** | **rs12410869** | **5.21E-13** | **1.000** | **ARNT** |
| 1q21.3 | ARNT | rs36008098 | 8.55E-13 | 0.02 | rs12410869 | 5.21E-13 | 1.000 | ARNT |
| **1q42.12** | **PARP1** | **rs9426568** | **3.53E-13** | **1.00** | **rs1858550** | **1.68E-13** | **0.996** | **PARP1** |
| **1q42.12** | **MIXL1** | **rs9426568** | **3.53E-13** | **0.57** | **rs1858550** | **1.68E-13** | **0.996** | **PARP1** |
| **1q42.12** | **MIXL1** | **rs1858550** | **1.68E-13** | **0.25** | **rs1858550** | **1.68E-13** | **1.000** | **PARP1** |
| 1q42.12 | ADCK3 | rs9426568 | 3.53E-13 | 0.01 | rs1858550 | 1.68E-13 | 0.996 | PARP1 |
| 1q42.12 | PSEN2 | rs9426568 | 3.53E-13 | 0.01 | rs1858550 | 1.68E-13 | 0.996 | PARP1 |
| **5p13.2** | **SLC45A2** | **rs250417** | **2.30E-12** | **0.09** | **rs250417** | **2.30E-12** | **1.000** | **SLC45A2** |
| **21q22.3** | **MX2** | **rs408825** | **3.21E-15** | **0.12** | **rs408825** | **3.21E-15** | **1.000** | **MX2** |
| **21q22.3** | **MX2** | **rs443099** | **3.50E-15** | **0.09** | **rs408825** | **3.21E-15** | **1.000** | **MX2** |
| 21q22.3 | BACE2 | rs364525 | 3.35E-15 | 0.04 | rs408825 | 3.21E-15 | 0.996 | MX2 |
| 21q22.3 | BACE2 | rs416981 | 3.28E-15 | 0.03 | rs408825 | 3.21E-15 | 1.000 | MX2 |
| 21q22.3 | BACE2 | rs408825 | 3.21E-15 | 0.02 | rs408825 | 3.21E-15 | 1.000 | MX2 |
| 21q22.3 | BACE2 | rs443099 | 3.50E-15 | 0.02 | rs408825 | 3.21E-15 | 1.000 | MX2 |
| **22q13.1** | **APOBEC3G** | **rs132941** | **1.61E-12** | **0.12** | **rs132941** | **1.61E-12** | **1.000** | **PLA2G6** |

\*eCAVIAR (Hormozdiari *et al. AJHG* 2016) was used for testing colocalization of melanocyte eQTL and melanoma GWAS signal. Fifty SNPs upstream and downstream of GWAS lead SNP in each locus were chosen to quantify the probability of the variant to be causal both in GWAS and eQTL studies
Colocalization posterior probability (CLPP): probability that the same variant is causal in both GWAS and eQTL. Only the genes of CLPP > 0.01 and SNPs in perfect LD ( $r^2 > 0.99$ in 1KG EUR population) with the GWAS lead SNP are presented. Genes of CLPP > 0.05 are shown in bold
Melanoma GWAS lead SNP: the lowest *P*-value SNP in the locus based on fixed effect model from Law *et al (Nat. Genet.* 2015) study
GWAS *P*-value: melanoma GWAS *P*-value (fixed model) of the SNP in the left column
$r^2$ : $r^2$ between the SNP from the eCAVIAR analysis and the melanoma GWAS lead SNP of the given locus (1000 Genomes, EUR)
Nearest expressed gene: gene whose gene body is closest to the melanoma GWAS SNP and is expressed in melanocytes at median TPM > 0

**Figure 4.**
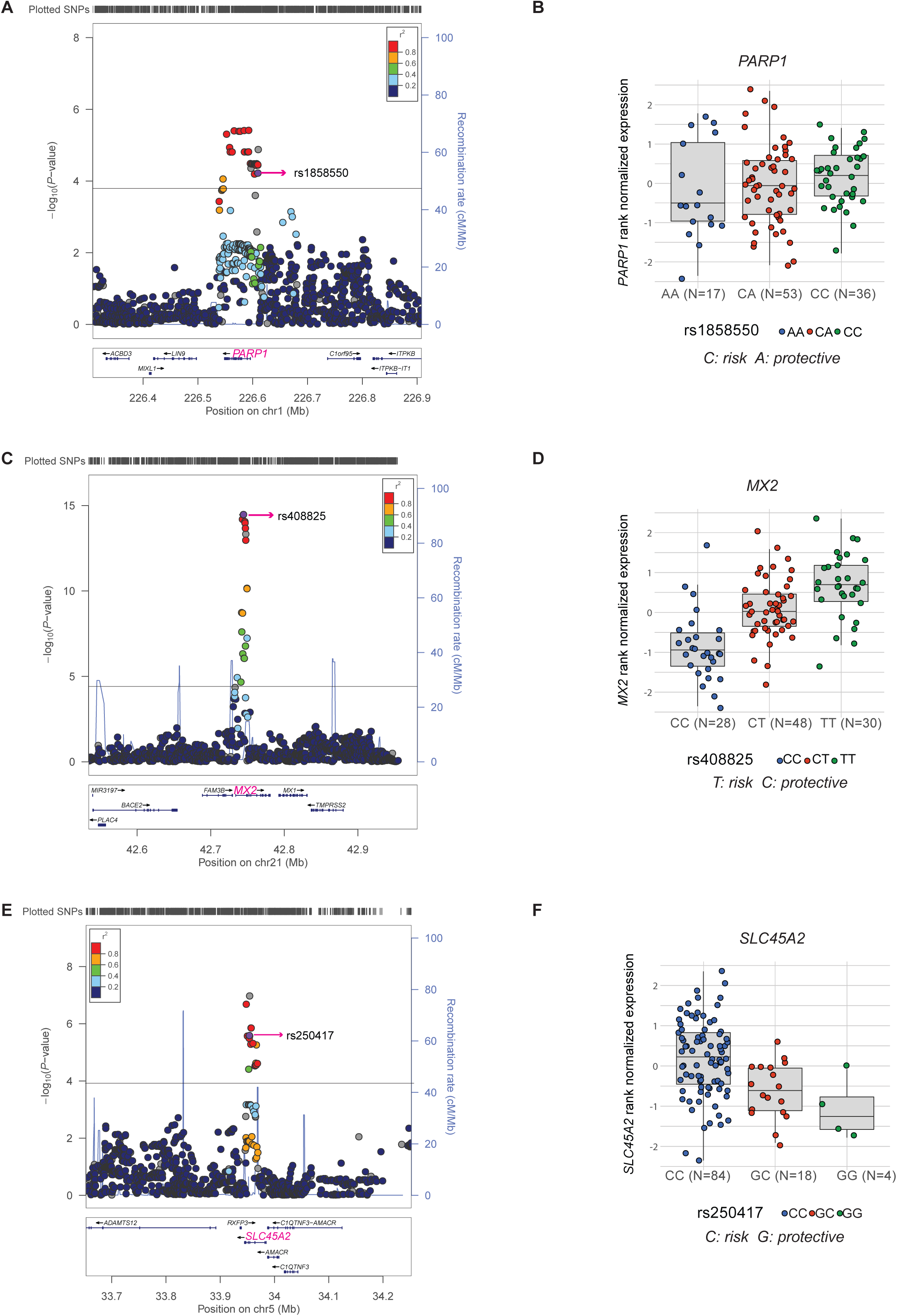
Melanoma GWAS signals colocalizing with melanocyte eQTL. (*A*, *C, E*) LocusZoom plots present the nominal eQTL *P*-values of all tested local SNPs in 300-400 kb windows for three significant eQTL genes from three melanoma GWAS loci: (*A*) *PARP1*; (*C*) *MX2*; and (*E*) *SLC45A2*. The gene being measured is highlighted in pink, the index melanoma risk SNP is labeled and highlighted in purple, and *r^2^* (based on 1000G EUR) of all other SNPs to the index SNP is color-coded. SNPs with missing LD information with the index SNP are shown in grey. Horizontal lines are shown for nominal *P*-value cutoff for significant eQTLs. Genomic coordinates are based on hg19. (*B, D, F*) Box plots present melanocyte expression differences of each gene in relation to the genotypes of the index SNP. Melanoma risk and protective alleles are shown for each locus.

We then performed permutation analyses to test for statistically significant enrichment of eQTLs from the four tissue types (including TCGA melanomas) in melanoma GWAS using four tiers of GWAS *P*-value thresholds (5 × 10^−5^, 5 × 10^−6^, 5 × 10^−7^, and 5 × 10^−8^; **Supplemental Table 17**). The results indicated that melanoma-associated SNPs using all four thresholds are significantly enriched (at least 2-fold) in eQTLs. Notably, the number of GWAS loci displaying true overlap was much higher (8-12 loci) for melanocyte eQTLs than for two types of skin tissue or melanoma tumors (2-7 loci).

### TWAS using melanocyte eQTL data identified four novel melanoma-associated loci

eQTL data can be utilized for transcriptome-wide association studies (TWAS) to impute gene expression levels into GWAS datasets. We performed a TWAS (Gusev et al. 2016) using summary statistics from the melanoma GWAS meta-analysis (Law et al. 2015) and the melanocyte eQTL dataset as the reference dataset (see **Methods**). Using 3,187 eGenes passing a conservative cutoff for heritability estimates (*P* < 0.01) (**Supplemental Table 18-19**), TWAS identified genes at three known melanoma loci at a genome-wide significant level (*MAFF* on Chr22q13.1, *CTSS* on Chr1q21.3, *CASP8* on Chr2q33-q34), with a fourth locus being suggestive (*PARP1* on Chr1q42.1) (Table 3). TWAS further identified novel associations with melanoma at four genomic loci at a genome-wide significant level (*ZFP90* at Chr16q22.1, *HEBP1* at Chr12p13.1, *MSC* and *RP11-383H13.1* at Chr8q13.3, and *CBWD1* at Chr9p24.3) (**Table 3** and **Fig. 5**).

**Table 3.** Top melanoma TWAS genes using melanocyte eQTL as a reference set.

| Gene Symbol | Chr | GWAS Best SNP | GWAS Z-score<sup>1</sup> | eQTL Best SNP | eQTL Z-score | GWAS Z-score<sup>2</sup> | # of SNP | # of Weight | Model | TWAS Z-score | TWAS P-value | FDR passed | GWAS locus |
| --- | --- | --- | --- | --- | --- | --- | --- | --- | --- | --- | --- | --- | --- |
| *MAFF* | 22 | rs132985 | -6.91 | rs738322 | -5.14 | -6.69 | 340 | 2 | enet | 6.67 | **2.60E-11** | Y | 22q13.1 |
| *CTSS* | 1 | rs12410869 | -7.22 | rs7521898 | 3.33 | -6.42 | 315 | 6 | enet | -6.32 | **2.65E-10** | Y | 1q21.3 |
| *ZFP90* | 16 | rs7184977 | 5.07 | rs11075688 | 1.06 | 3.96 | 319 | 319 | blup | 5.20 | **1.95E-07** | Y | New |
| *HEBP1* | 12 | rs2111398 | 5.08 | rs1684387 | -5.32 | 5.08 | 605 | 5 | lasso | -5.05 | **4.48E-07** | Y | New |
| *CASP8* | 2 | rs10931936 | 5.50 | rs3769823 | -3.71 | 4.45 | 372 | 3 | lasso | -4.61 | **4.06E-06** | Y | 2q33-q34 |
| *MSC* | 8 | rs1481853 | -4.65 | rs6983160 | -3.83 | -4.60 | 553 | 2 | lasso | 4.60 | **4.27E-06** | Y | New |
| *CBWD1* | 9 | rs661356 | 4.79 | rs2992854 | 4.99 | -1.96 | 462 | 1 | lasso | -4.54 | **5.52E-06** | Y | New |
| *RP11-383H13.1* | 8 | rs1481853 | -4.65 | rs6983160 | -2.75 | -4.60 | 693 | 4 | lasso | 4.50 | **6.68E-06** | Y | New |
| *GPRC5A* | 12 | rs2111398 | 5.08 | rs1684387 | -4.02 | 5.08 | 584 | 11 | enet | -4.19 | 2.80E-05 | N | New |
| *RBBP5* | 1 | rs11240466 | 4.09 | rs10900456 | 1.41 | 3.98 | 623 | 623 | blup | 3.59 | 3.37E-04 | N | New |
| *ATP6V1G2-DDX39B* | 6 | rs2239704 | 4.55 | rs2523504 | 3.51 | 3.54 | 278 | 1 | lasso | 3.54 | 3.97E-04 | N | New |
| *CDH1* | 16 | rs7184977 | 5.07 | rs4076177 | 2.71 | 4.52 | 345 | 345 | blup | 3.47 | 5.27E-04 | N | New |
| *CHCHD6* | 3 | rs9851451 | 3.46 | rs9822602 | -2.73 | 3.44 | 635 | 3 | lasso | -3.44 | 5.78E-04 | N | New |
| *CTD-2003C8.2* | 11 | rs1554519 | 4.51 | rs7932891 | 5.66 | 3.32 | 746 | 4 | lasso | 3.43 | 6.08E-04 | N | New |
| *UQCC1* | 20 | rs2425025 | 6.95 | rs6060369 | -3.14 | -3.00 | 353 | 353 | blup | 3.42 | 6.21E-04 | N | New |
| *PLXNA1* | 3 | rs9851451 | 3.46 | rs4679317 | -2.20 | 3.42 | 484 | 2 | lasso | -3.42 | 6.33E-04 | N | New |
| *PARP1* | 1 | rs1865222 | -7.12 | rs3219090 | -1.30 | -6.86 | 404 | 404 | blup | 3.41 | 6.57E-04 | N | 1q42.12 |

Chr: chromosome
GWAS Best SNP: rsID of the most significant melanoma GWAS SNP for the TWAS gene after QC and IMPG imputation by FUSION program
GWAS Z-score<sup>1</sup>: GWAS Z-score of the most significant GWAS SNP in the locus
eQTL Best SNP: rsID of the best eQTL SNP in the locus
eQTL Z-score: eQTL Z-score of the best eQTL SNP in the locus
GWAS Z-score<sup>2</sup>: GWAS Z-score for the best eQTL SNP

**Figure 5.**
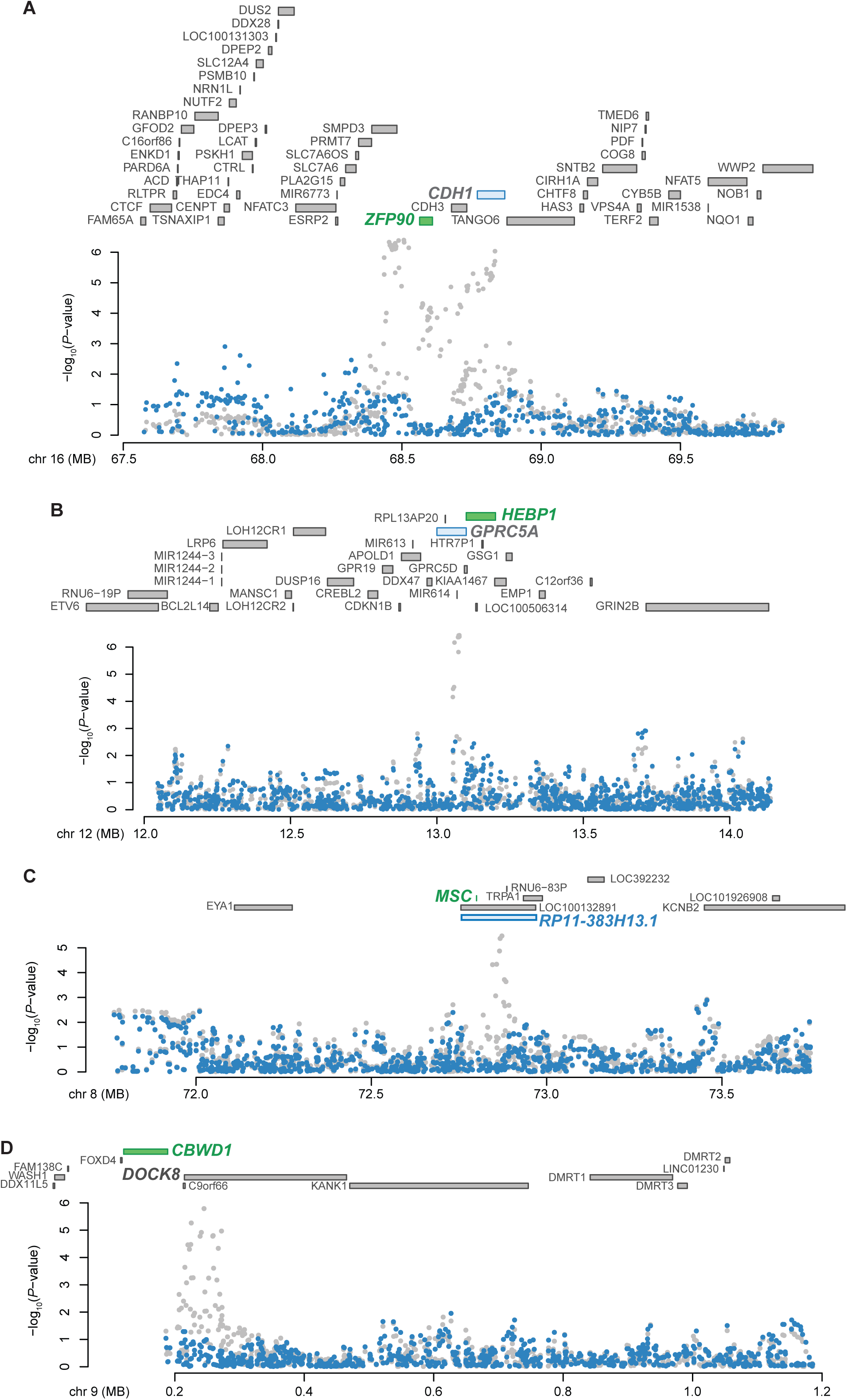
TWAS using melanocyte eQTL data as a reference set identified five new melanoma-associated genes in four new loci. (*A*) The new melanoma TWAS gene, *ZFP90* on Chromosome 16 (TWAS *P* = 1.95 × 10^−7^, TWAS Z = 5.2) is shown in green, along with a second marginally significant gene, *CDH1* (*P* = 5.27 ×10^−4^, Z = 3.47) and other annotated genes at the locus (coordinates are hg19). The Manhattan plot presents the melanoma GWAS *P*-values before (gray) and after (blue) conditioning on imputed melanocyte-specific gene expression of the gene in green, (*ZFP90* in this locus). (*B*) A similar plot of for the melanoma TWAS gene *HEBP1* (TWAS *P* = 4.65 × 10^−7^, TWAS Z=−5.04) and a second marginally significant gene, *GPRC5A* (TWAS *P*= 2.8 × 10^−5^, TWAS Z = −4.19) on Chromosome band 12p13.1. (C) A similar plot for two new melanoma TWAS genes, *MSC* (*P* = 4.27 × 10^−6^, Z = 4.6) and *RP11-383H13.1* (*P* = 6.68 × 10^−6^, Z = 4.5) on Chromosome band 8q13.3. The Manhattan plot shows the melanoma GWAS *P*-values before (gray) and after (blue) conditioning on imputed melanocyte-specific gene expression of *MSC*. (*D*) A similar plot of new melanoma TWAS gene, *CBWD1* (*P* = 5.52 × 10^−6^, Z = −4.54) and a marginally significant gene, *DOCK8* (*P* = 2.7 × 10^−3^, Z = 2.99) on Chromosome band 9p24.3.

We additionally performed TWAS using each of the 44 GTEx tissue types as reference eQTL datasets. Forty-three GTEx tissue types identified one or more melanoma TWAS genes at a genome-wide significant level with a median of three genes per dataset. Tibial nerve tissue identified the largest number of genes (11 genes) while melanocytes ranked third (**Supplemental Table 19**). Collectively, melanocyte and GTEx datasets identified 22 TWAS genes at six previously known melanoma GWAS loci (Chr1q21.3, Chr1q42.1, Chr2q33-q34, Chr15q13.1, Chr21q22.3, and Chr22q13.1) as well as nine TWAS genes at eight novel loci. Melanocyte eQTLs alone identified five of the nine novel TWAS genes, three of which (*HEBP1, MSC* and *RP11-383H13.1*) were only genome-wide significant when using the melanocyte eQTL dataset (**Supplemental Table 20**). In contrast, none of the 44 GTEx tissue datasets produced more than one novel association for melanoma. Five novel melanoma TWAS genes added from 44 GTEx tissue types are *ERCC2* on Chr19q13.32, *KIF9* on Chr3p21.31, *MRAP2* on Chr6q14.2, and *ZBTB4* on Chr17p13.1. Finally, we conducted conditional analyses on the TWAS loci displaying marginally significant associations with multiple genes from melanocyte and GTEx tissue datasets. The analyses identified 15 jointly significant genes from 14 loci (**Supplemental Table 20; Supplemental Material**), including *CTSS* from the multi-gene locus at Chr1q21.3.

## DISCUSSION

In this study, we established a cell-type specific eQTL dataset using primary cultures of human melanocytes. Our dataset identified a unique set of *cis-* and *trans-*eQTLs that are distinct from eQTLs of skin tissues. Melanocyte eQTLs are enriched in melanocyte-specific *cis-*regulatory elements and considerably improved melanoma GWAS annotation. Using this dataset, we further identified novel melanoma TWAS loci. Our data highlight the utility of building even a modestly sized cell-type specific dataset.

Over a third of melanocyte eGenes were unique to melanocytes and not present in skin tissue datasets. GO analyses suggested that genes directly involved in melanin synthesis as well as those in lysosome and metabolic pathways were enriched in melanocyte eGenes among others. These observations are consistent with broad-based pleiotropic cell functions for genes expressed in melanocytes including lysosome-related functions of melanin synthesis and transfer process (Sitaram and Marks 2012). Our dataset was built with newborn males of primarily European descent aiming to align with the most relevant population for melanoma incidence. As there are gender differences observed in melanoma risk and mortality among others (US National Cancer Institute’s Surveillance Epidemiology and End Results Database; 2012-2014) (Scoggins et al. 2006; Wendt et al. 2018), the current male-only dataset cannot address gender-specific risk and related questions, which warrants future study.

Through *trans*-eQTL analysis, the melanocyte dataset identified IRF4, or Interferon regulatory factor 4, as a potential regulator of melanocytic-lineage specific gene expression for a set of downstream genes. *trans*-eQTL*s* were shown to be more cell-type specific than *cis-*eQTLs, and cell composition heterogeneity was proposed as a potential reason for low number of *trans*-eQTL*s* observed in bulk tissue data (Westra and Franke 2014; The GTEx Consortium et al. 2017), suggesting that our single-cell type dataset might have facilitated the identification of the IRF4 *trans*-eQTL network in melanocytes. rs12203592 is a *cis*-eQTL in several other GTEx tissue types including whole blood, perhaps reflecting a better known function of *IRF4* in immune responses (Huber and Lohoff 2014). However, IRF4 has a documented role in melanocyte development, regulating expression of an enzyme essential in the production of melanin, tyrosinase (Praetorius et al. 2013). Given that IRF4 appears to function in regulation of distinct cell type-specific processes, four rs12203592 *trans*-eQTL genes identified in melanocytes, including an Interferon-Stimulated Gene, *TMEM140* (Kane et al. 2016), as well as perhaps a considerably larger subset of marginal *trans*-eQTL genes, could be good candidates for direct targets of IRF4. Further experimental assessment of IRF4 binding on the genomic regions of these *trans*-genes will provide additional support of this finding.

Through colocalization and TWAS, melanocyte eQTL identified unique candidate melanoma susceptibility genes for some known loci and also corroborated other datasets including skin, in identifying candidate genes for other loci. Three melanoma loci displaying colocalization with melanocyte eQTL were also supported by TWAS from one or more eQTL datasets (*PARP1, ARNT*, and *MX2*). On the other hand, some loci with larger LD blocks displayed variability in target gene prediction across different datasets as well as between eCAVIAR and TWAS approaches. For the melanoma locus at Chr1q21.3, a total of eight genes were colocalized from three eQTL datasets, and TWAS nominated nine genes. Each top CLPP score gene from melanocyte and skin datasets (*ARNT, CERS2*, and *SETDB1*) were also supported by TWAS in more than one tissue type, while TWAS joint/conditional analyses identified *CTSS* as the major signal of this locus. These data imply that multiple statistical approaches using diverse tissue types including the cell-type of disease origin is beneficial to robust target gene prediction. Collaboration of single cell type and whole tissue eQTL was also exemplified in *ASIP* for hair color GWAS colocalization (**Supplemental Material**).

TWAS also identified novel melanoma loci by leveraging tissue-specific eQTL datasets and reducing multiple testing burden associated with GWAS. While identification of trait-associated gene expression differences via TWAS cannot be taken to imply causality for a specific gene, TWAS may nonetheless nominate plausible candidate risk genes at significant loci. Here, we identified five genes at four new melanoma susceptibility loci (*ZFP90* on Chr16q22.1, *HEBP1* on Chr12p13.1, *MSC* and *RP11-383H13.1* on Chr8q13.3, and *CBWD1* on Chr9p24.3) using melanocyte eQTL as a reference set, and four additional new genes/loci (*ERCC2, KIF9, MRAP2*, and *ZBTB4*) using 44 GTEx tissue types.

While most of these genes have known functions that might have relevance in melanomagenesis (**Supplemental Material**), *ERCC2* on Chr19q13.32, among them, is a nucleotide excision repair gene targeting UV-induced DNA damage and implicated in Xeroderma Pigmentosum (Taylor et al. 1997). *MRAP2* on Chr6q14.2 encodes melanocortin-2-receptor accessory protein 2, which interacts with all melanocortin receptor proteins (MCRs) together with MRAP1 to regulate cell surface expression of MCRs (Ramachandrappa et al. 2013). Seemingly relevant functions of these new candidate genes warrant further studies on their roles in melanomagenesis. In all, our primary melanocyte eQTL dataset considerably advanced identification of candidate melanoma susceptibility genes from known and new melanoma loci through multiple approaches, which highlights the unique value of cell-type specific eQTL datasets.

## METHODS

### Melanocyte culture

We obtained frozen aliquots of melanocytes isolated from foreskin of 106 healthy newborn males who are mainly of European descent following an established protocol (Halaban et al. 2000) from the SPORE in Skin Cancer Specimen Resource Core at Yale University. Cells were grown in Dermal Cell Basal Medium (ATCC^®^ PCS-200-030™) supplemented with Melanocyte Growth Kit (ATCC^®^ PCS-200-041™) and 1% Amphotericin B/Penicillin/Streptomycin (120-096-711, Quality Biological) at 37°C with 5% CO_2_, and trypsinized with Trypsin/EDTA Solution (R-001-100, Cascade Biologics) as well as Trypsin Neutralizer Solution (R-002-100, Cascade Biologics). Media was changed every 2-3 days when necessary. Throughout the whole process, two specific lot numbers of medium and supplement were used for consistency, and for the final passage of at least 2 days before harvesting cells for RNA and DNA, a single lot of medium and supplement was used for the whole panel. Every step of DNA/RNA isolation, and sequencing/genotyping processes were also performed in re-randomized batches. Before harvesting the cells, media was taken and tested for mycoplasma contamination using MycoAlert PLUS mycoplasma detection kit (LT07-710, Lonza). All 106 samples were negative for mycoplasma contamination.

### Genotyping and imputation

Genomic DNA was isolated from frozen pellets in randomized batches using the Gentra Puregene Cell Kit (158745, Qiagen). After DNA quantity and quality assessment, DNA samples were genotyped on the Illumina OmniExpress arrays (HumanOmniExpress-24-v1-1-a) in randomized batches of 24 samples per chip at the Cancer Genomics Research Laboratory of the Division of Cancer Epidemiology and Genetics (NCI/NIH). After genotype quality control, genotypes were imputed using Michigan Imputation Server (Das et al. 2016) based on 1000 Genomes (Phase 3, v5) reference panel (1000 Genomes Project et al. 2015) and Mixed population, and using SHAPEIT for pre-phasing. Post-imputation genetic variants (single nucleotide variants (SNP) and small insertion-deletion (indel) polymorphisms) with MAF<0.01 or imputation quality scores (R-squared) <0.3 were removed from the final analysis. Overall, ~713,000 genotypes were obtained, and 10,718,646 genotypes were further imputed. Due to the small sample size, we included all samples that passed genotyping QC but histologically carry a range of African and Asian ancestry measured by ADMIXTURE analysis, while accounting for ancestry in the further analyses as covariates. For eQTL analysis, we included the top 3 genotyping principal components as covariates. The principal components analysis for population substructure was performed using the *struct.pca* module of GLU (Wolpin et al. 2014), which is similar to EIGENSTRAT(Price et al. 2006).

### RNA sequencing and data processing

Cells were harvested at log phase by washing with cold PBS on ice followed by lysis with QIAzol lysis reagent and stored at −80°C. Total RNA was isolated using miRNeasy Mini Kit (217004, Qiagen) in randomized batches. RNA quantity and quality was assessed using a NanoDrop8000 spectrophotometer and Bioanalyzer, which yielded RIN>9 for all 106 samples and RIN=10 for >75% of the samples. Poly-A selected stranded mRNA libraries were constructed from 1 μg total RNA using Illumina TruSeq Stranded mRNA Sample Prep Kits according to manufacturer’s instructions except where noted. The pooled (barcoded and randomized) libraries were sequenced on multiple lanes of a HiSeq2500 using version 4 chemistry to achieve a minimum of 45 million 126 base paired reads (average of ~87.9 million reads). STAR (version 2.5.0b) (Dobin et al. 2013) was used for aligning reads to the human genomic reference (hg19) with the gene annotation from GENCODE Release 19 (https://www.gencodegenes.org/releases/19.html). VerifyBamID was used to check whether the reads were contaminated as a mixture of two samples by and no contamination was found (Jun et al. 2012). RSEM (version 1.2.31, http://deweylab.github.io/RSEM/) was used to quantify the gene expression followed by the quantile normalization. For eQTL analysis, we used the same method of post-processing gene expression data as the GTEx project (http://www.gtexportal.org). Genes were selected based on expression thresholds of >0.5 RSEM in at least 10 samples and ≥6 reads in at least 10 samples. After processing, 19,608 genes were expressed above cutoff levels in primary melanocytes. For each gene, expression values were further inverse quantile normalized to a standard normal distribution across samples. To control for hidden batch effects and other confounding effects that could be reflected in the expression data, a set of covariates identified using the Probabilistic Estimation of Expression Residuals (PEER) method (Stegle et al. 2010) was calculated for the normalized expression matrices. The top 15 PEER factors were determined based on the sample size and optimizing for the number of eGenes discovered (15 factors for N<150).

### Identification of *cis*-eQTLs in primary melanocytes

*Cis*-eQTL analysis was performed closely following a recent standard procedure adopted by GTEx (Aguet et al. 2016). In brief, *cis*-eQTL mapping was performed using FastQTL (Ongen et al. 2016), using the expression data, imputed genotype data, and covariates described above. First, nominal *P*-values were generated for each variant-gene pair by testing the alternative hypothesis that the slope of a linear regression model between genotype and expression deviates from 0. Genetic variants located within +/− 1Mb of the TSSs for each gene were tested for *cis*-eQTL effects of the corresponding gene. Variants in imputed VCF were selected based on the following thresholds: the minor allele was observed in at least 10 samples (the minor allele frequency was ≥ 0.05). Second, the beta distribution-adjusted empirical *P*-values from FastQTL were used to calculate q-values (Storey and Tibshirani 2003) and a false discovery rate (FDR) threshold of ≤0.05 was applied to identify genes with a significant eQTL (“eGenes”). The adaptive permutations mode was used with the setting “--permute 1000 10000”. The effect size of the eQTLs was defined as the slope of the linear regression and is computed as the effect of the alternative allele (ALT) relative to the reference allele (REF). Last, to identify the list of all significant variant-gene pairs associated with eGenes, a genome-wide empirical *P*-value threshold, *p_t_*, was defined as the empirical p-value of the gene closest to the 0.05 FDR threshold. *p_t_* was then used to calculate a nominal p-value threshold for each gene based on the beta distribution model (from FastQTL) of the minimum *P*-value distribution *f*(*p*_min_) obtained from the permutations for the gene. Specifically, the nominal threshold was calculated as *F^−1^*(*p_t_*), where *F^−1^* is the inverse cumulative distribution. For each gene, variants with a nominal p-value below the gene-level threshold were considered significant and included in the final list of variant-gene pairs. The number of identified eGenes and significant eQTLs was approximately three times higher than those from data analyzed without using PEER factors as covariates (**Supplemental Table 2**). Application of PEER factors almost doubled the number of eGenes known to be related to pigmentation phenotypes (0.8% vs 1.5%; Fisher exact test *P*-value = 0.0335).

### Pairwise eQTL sharing between primary melanocytes and 44 GTEx tissues

To assess replication of *cis*-eQTL and eGenes in the publicly available Genotype Tissue Expression (GTEx) project (The GTEx Consortium 2013; Aguet et al. 2016), we collected eQTL results from 44 tissue types with >= 70 samples (The GTEx Analysis V6p). The eGene and significant SNP-gene associations based on permutations were collected (GTEx_Analysis_v6p_eQTL.tar) and every SNP-gene association test (including non-significant tests) were download from GTEx website. To test the sharing of all significant SNP-gene pairs of our melanocytes eQTL study with the ones identified in 44 tissue types by GTEx, we used the threshold of *FDR* <0.05 and calculated the pairwise *π*_1_ statistics. We used Storey’s QVALUE software (Storey and Tibshirani 2003) (https://github.com/StoreyLab/qvalue) to calculate the *π*_1_, which indicates the proportion of true positives. A heat map was drawn based on the pairwise *π*_1_ values. The pairwise *π*_1_ statistics are reported for single-tissue eQTL discoveries in each tissue. Higher *π*_1_ values indicate an increased replication of eQTLs. Tissues are grouped using hierarchical clustering on rows and columns separately with a distance metric of 1 – ρ, where ρ is the Spearman correlation of *π*_1_ values. *π*_1_ is only calculated when the gene is expressed and testable in both the discovery and the replication tissues.

### Identification of *trans*-eQTL*s* in primary melanocytes

*trans*-eQTL analysis was performed for SNPs that are located over 5Mb away from the TSS of each gene or on a different chromosome. Genes of mappability < 0.8 or overlapping low complexity regions defined by RepeatMasker library were excluded from the analysis. The nominal *P*-values for gene-SNP pairs in *trans*-eQTL analysis were calculated using the Matrix-eQTL program (Shabalin 2012). We performed multiple testing to identify significant *trans*-eQTLs following our previous approach (Shi et al. 2014). SNPs with call rates <0.9 or minor allele frequencies (MAF) <0.05 were excluded, as were SNPs out of Hardy Weinberg equilibrium with p<10^−6^. For each nominal *P*-value threshold *p*, we calculated the number of genes (denoted as *N*_1_(*p*)) that has at least one SNP in its trans region with nominal *P*-value less than the threshold *p*. Here, *N*_1_(*p*) denotes the number of *trans*-eQTL genes at *P*-value threshold *p*. Next, we performed 100 permutations to estimate the number of genes (denoted as *N*_0_(*p*)) detected to have *trans*-eQTL signals at nominal *P*-value *p* under the global null hypothesis. By definition, one can calculate FDR as *FDR = N*_0_(*p*)*/N*_1_(*p*). We chose *p* = 3.25 × 10^−11^ to control FDR at a desired level of 0.1.

### Identifying *cis*-mediators for *trans*-eQTLs in primary melanocytes

We applied the Genomic Mediation analysis with Adaptive Confounding adjustment (GMAC) (Yang et al. 2017) algorithm to identify *cis*-mediators for *trans*-eQTLs in primary melanocytes eQTL data. Only the trios with evidence of both *cis* and *trans* association were kept. The *cis-*eSNP with smallest *P*-value for each gene (eQTL FDR<0.05) and trans-association *P*-value is less than 10^−5^ was selected as one trio. Up to 5 PEER factors and other covariates (top 10 genotype PCs) were adjusted. 100,000 permutations for testing mediation were performed and trios with suggestive mediation were reported using mediation *P*-value threshold <0.05.

### Allele-specific Expression (ASE)

ASE analysis was performed based on the GATK best practices pipeline in allelic expression analysis published by the Broad Institute (Castel et al. 2015). We included heterozygous loci in exonic regions when the imputation quality was R^2^>0.9 and probability of heterozygosity >95%. After the quality control, we evaluated the significance of allelic imbalance using a binomial test in each individual level, comparing the observed to the subject- and genotype-specific expected allele ratios (Ongen et al. 2014). To minimize false ASE events resulting from mapping bias specific to different DNA bases, we compared the observed allelic ratio for each coding heterozygous variant to the overall ratio for that specific allele pair in each sample (i.e. for each of the following pairs: AC, AG, AT, CA, CG, CT, GA, GC, GT, TA, TC, TG heterozygotes) (Lappalainen et al. 2013), so that we were able to take into account the expected allele imbalance in the ASE analysis. In addition, the effect size of allelic expression (AE, defined as |0.5-Reference ratio|) were calculated. We defined significant ASE genes as genes with at least one genetic variant exhibiting a minimum effect size of 0.15 or a significant difference from the expected allele ratio of 0.5 at FDR <0.05 (calculated using the Benjamini and Hochberg approach) (Benjamini and Hochberg 1995) in one or more individuals. Significant ASE genes were then grouped into melanocyte eGenes and non-eGenes, and |Mean AE| values as well as percentage of individuals displaying allelic imbalance were compared between two groups (Wilcoxon Rank Sum and Singed Ranked Test).

### Assessing enrichment in putative functional elements

To assess the enrichment of *cis*-eQTL in putative functional elements of primary melanocytes, we collected the DNase-seq and ChIP-seq data from the Epigenome Roadmap Project (http://www.roadmapepigenomics.org) (Roadmap Epigenomics et al. 2015). For each putative functional element, we merged peak callings from all samples into one, and all the significant melanocyte eQTL SNP-Gene pairs were used for the enrichment analyses using a similar method to a recent publication (Zhang et al. 2017). Briefly, we performed randomizations for testing whether an eQTL SNP set is enriched for given histone mark regions. Note that the following procedure controls for the distribution of minor allele frequencies of a given eQTL SNP set: 1.) For *K* eQTL SNPs, we determined the number (denoted as *X*_0_) of eQTL SNPs functionally related with the histone mark, 2.) We randomly sampled 10,000 SNP sets. Each SNP set had *K* SNPs in linkage equilibrium, with minor allele frequency distribution similar to the original *K* eQTL SNPs. For the *n^th^* sampled SNP set, we calculated the number (denoted as *x_n_*) of SNPs functionally related with the histone mark. We had {*x*_1_,…, *x*_10000_}, corresponding to the sampled 10000 SNP sets, and 3.) Enrichment fold change was calculated as 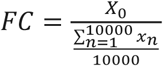, where the denominator represented the average number of SNPs functionally related with the histone mark under the null hypothesis. The *P*-value for enrichment was calculated as *P =* {*n: x_n_ ≥* X_0_}/10000, i.e., the proportion of SNP sets functionally more related with the histone mark than the given eQTL SNP set. If *x_n_ < X*_0_ for all sample SNP set, we reported P value as *P <* 10^−4^. In addition, we also assessed enrichment of *cis*-eQTLs in different genomic regions including 5’/3’-UTR, promoter, exon, intron, intergenic and lncRNA region as described in R annotatr package (https://github.com/hhabra/annotatr).

### Enrichment of melanoma GWAS variants in eQTLs

Two methods were used to evaluate if the melanoma GWAS variants were enriched in eQTLs of different datasets. First, QQ plots were used to show the differences in melanoma association *P*-values between the significant eQTL SNPs and non-eQTL SNPs. For all the GWAS variants, we first performed LD pruning using PLINK (r^2^ = 0.1 and window size 500 kb), so that the remaining SNPs were independent. Then, based on eQTL data, these pruned SNPs were classified into two groups. If a SNP is an eQTL or is in LD (r^2^ > 0.8) with an eQTL SNP, the SNP is classified as eQTL SNPs. Otherwise, it is classified as non-eQTL SNPs. QQ plots were generated using the melanoma GWAS *P*-values from the most recent meta-analysis (Law et al. 2015) for eQTL SNPs versus non-eQTL SNPs. Deviation from the 45-degree line indicates that melanoma GWAS SNPs are strongly enriched in eQTL SNPs. The lambda values were estimated using the “estlambda2” function in R package “QQperm”. For the second method, a similar simulation procedure was applied to identify overlap and test for enrichment of eQTLs in melanoma GWAS SNPs (Hannon et al. 2016). All significant eQTLs were ‘clumped’ based on the best eQTL *p* value using PLINK to create a list of quasi-independent SNPs (r^2^<0.25 for all pairs of SNPs within 250kb) and to prevent LD between SNPs in the set biasing the results. A more stringent clumping procedure was used for SNPs located in Chr5:25000000-35000000, where the window for pairwise SNP comparisons was extended to 10,000kb. 1,000,000 simulated sets matched for allele frequency were drawn to calculate the expected overlap between the eQTL SNP and melanoma GWAS variants at four GWAS significance thresholds (*P*<5E-5, 5E-6, 5E-7, 5E-8) and generate empirical *P*-values. Empirical significance for enrichment of eQTLs in GWAS variants was ascertained by counting the number of simulations with at least as many SNP sets and dividing by the number of simulations performed. Fold change statistics were calculated as the true overlap divided by the mean overlap of these simulations.

### Melanoma heritability enrichment of tissue-specific genes

We used stratified LD score regression implemented in LDSC program (https://github.com/bulik/ldsc) to estimate the enrichment of melanoma heritability for SNPs around tissue- and cell-type specific genes as described previously (Finucane et al. 2018). We downloaded the gene expression file from GTEx Portal (GTEx Analysis V6p Gene RPKM file: GTEx_Analysis_v6p_RNA-seq_RNA-SeQCv1.1.8_gene_rpkm.gct.gz) and quantified our melanocyte RNA-Seq data as RPKM using the same method (RNA-SeQCv1.18) (DeLuca et al. 2012). To reduce batch effects, quantile normalization was applied to combined melanocyte and GTEx RNA-Seq RPKM values. For GTEx data, we used the ‘SMTSD’ variable (‘Tissue Type, more specific detail of tissue type’) to define our tissues and the ‘SMTS’ variable (‘Tissue Type, area from which the tissue sample was taken’) to define the tissue categories for t-statistic computation (see **Supplemental Table 8** for the tissue categories). We treated the tissue category for melanocytes as “Skin”. To define the tissue-specific genes, we calculated the t-statistic of each gene for a given tissue, excluding all samples from the same tissue category. For example, for melanocytes, we compared expression levels of each gene in the melanocyte samples to those of all other tissue samples in non-”Skin” categories to obtain a t-statistic. We selected the top 1,000, 2,000, and 4,000 tissue-specific genes by t-statistic, added a 100-kb window around their transcribed regions to define tissue-specific genome annotation, and applied stratified LD score regression on a joint SNP annotation to estimate the heritability enrichment against the melanoma GWAS meta-analysis (Law et al. 2015). The results using the top 4,000 tissue-specific genes showed significant enrichment (FDR < 0.05) for melanocyte and all three tissue types in the “Skin” category. The overall pattern was consistently observed in results using 2,000 and 1,000 genes, while melanocyte was significant in results from 2,000 but not in those from 1,000 genes (**Supplemental Table 8; Supplemental Figure 7**). Importantly, some of the top enriched tissues outside of the “Skin” category (e.g. Colon_Transverse) displayed high median expression level correlation with melanocytes (Pearson *r* = 0.95 between melanocyte and Colon_Transverse; data not shown).

### Colocalization analysis of GWAS and eQTL data

We performed colocalization analysis for 20 GWAS loci from the most recent GWAS meta-analysis using CAusal Variants Identification in Associated Regions (eCAVIAR, http://genetics.cs.ucla.edu/caviar/index.html). eCAVIAR is a statistical framework that quantifies the probability of the variant to be causal both in GWAS and eQTL studies, while allowing an arbitrary number of causal variants. For each locus, both GWAS and eQTL (from human melanocytes cultures in our study and two GTEx skin tissues) summary statistics of selected variants in that locus were extracted as the input for eCAVIAR. We selected 50 SNPs both upstream and downstream of the GWAS lead SNP for each GWAS locus. We computed the CLPP score with maximum number of two causal SNPs in each locus. We used CLPP >1% (0.01) cutoff for co-localization. Thus, for a given GWAS variant (either the lead SNP itself or the SNPs in near perfect LD with the lead SNP using the cutoff *r*^2^ > 0.99), an eGene with a CLPP score above the colocalization cutoff is considered a target gene. We also highlight the eGenes with CLPP > 0.05 as they were more robust across minor changes in analyses criteria compared to those on the borderline (between 0.01 and 0.05) in our analyses. For eCAVIAR analyses of nevus count GWAS, summary statistics for SNPs surrounding four previously-published nevus-only genome-wide significant loci were obtained from Duffy et al study (Biorxv, https://doi.org/10.1101/173112). GWAS summary statistics for 50 SNPs upstream and downstream the lowest *P*-value SNPs (rs12203592 for the locus at Chr6p25.3, rs869330 for Chr9p21.3, rs10521087 for Chr9q31.1-2, and rs4380 for Chr22q13.1) were extracted for the analysis. For pigmentation trait GWAS, GWAS studies were selected from the GWAS catalog using the keyword “pigmentation”, and reported lead SNPs were grouped based on the cytoband. The boundary of each region was set based on the union of the lead SNPs in the same cytoband, and 1Mb was added to each side of the boundary to look for the lowest *P*-value SNPs in the same region from the UK Biobank (UKBB) dataset. Three pigmentation traits available in UKBB dataset (skin pigmentation, ease of tanning, and hair color coded in a continuous scale of red, blonde, light brown, dark brown, and black) were chosen to obtain the GWAS summary statistics.

### Performing TWAS with GWAS summary statistics

We performed 45 transcriptome-wide association studies (TWAS) by predicting the function/molecular phenotypes into GWAS using melanoma GWAS summary statistics and both GTEx and melanocyte RNA-seq expression data. The new framework TWAS/FUSION (http://gusevlab.org/projects/fusion/) was used to perform the TWAS analysis, allowing for multiple prediction models, independent reference LD, additional feature statistics and cross-validation results (Gusev et al. 2016). In brief, we collected the summary statistics data including no significance thresholding in LD-score format (https://github.com/bulik/ldsc/wiki/Summary-Statistics-File-Format) from the most recently published cutaneous melanoma meta-analysis (Law et al. 2015). The precomputed expression reference weights for GTEx gene expression (V6) RNA-seq across 44 post mortem tissue were downloaded (http://gusevlab.org/projects/fusion/). We computed our functional weights from our melanocyte RNA-seq data one gene at a time. Genes that failed quality control during heritability check (using minimum heritability *P*-value 0.01) were excluded from the further analyses. We restricted the *cis*-locus to 500kb on either side of the gene boundary. A genome-wide significance cutoff (TWAS *P*-value < 0.05/number of genes tested) was applied to the final TWAS result. Multiple associated features in a locus were observed, and thus we performed the joint/conditional analysis to identify which are conditionally independent for each melanoma susceptibility locus using a permutation test with a maximum of 100,000 permutations and initiate permutation *P*-value threshold 0.05 for each feature. We also checked how much GWAS signal remained after conditioning on imputed expression levels of each associated feature by using “FUSION.post_process.R” script.

### Other analyses

*IRF4* motif enrichment analysis were performed using the AME module in The MEME Suite (http://meme-suite.org) and inputted shuffled sequences as control. *IRF4* motif were download from HOCOMOCO v10 database (http://hocomoco.autosome.ru/motif/IRF4_HUMAN.HI0MO.C). All the statistical analyses were performed in R (https://www.R-project.org/) (R Core Team 2018).

## ACKNOWLEDGMENTS

This work utilized the computational resources of the NIH high-performance computational capabilities Biowulf cluster (http://hpc.nih.gov). We acknowledge contributions to human melanocyte genotyping from The National Cancer Institute Cancer Genomics Research Laboratory (CGR) and to RNA sequencing from The NIH Intramural Sequencing Center (NISC), and NCI Center for Cancer Research Sequencing Facility (CCR-SF) and the Yale University Skin SPORE Specimen Resource Core. SM is supported by an Australian Research Council Fellowship. We thank A. Vu, L. Mehl, and H. Kong for proofreading the manuscript. This work has been supported by the Intramural Research Program (IRP) of the Division of Cancer Epidemiology and Genetics, National Cancer Institute, US National Institutes of Health.

The content of this publication does not necessarily reflect the views or policies of the US Department of Health and Human Services, nor does mention of trade names, commercial products, or organizations imply endorsement by the US government.

## DISCLOSURE DECLARATION

We do not have any conflict of interest to declare.

**Reference Genome Build Statement**

Our data was mapped to GRCh37/hg19 to allow maximum comparability with the GTEx and other public datasets we used in the manuscript. Mapping the reads of our data to the most current GRCh38 would not significantly affect the global eQTL analyses and conclusions of the current paper and only minor differences are expected.

